# From Armament to Ornament: Performance Trade-Offs in the Sexual Weaponry of Neotropical Electric Fishes

**DOI:** 10.1101/247072

**Authors:** Kory M. Evans, Maxwell J. Bernt, Matthew A. Kolmann, Kassandra L. Ford, James S. Albert

**Affiliations:** University of Louisiana at Lafayette, Department of Biology, P.O. Box 42451, Lafayette, LA 70504, USA, (MJB), (KLF), (JSA).; University of Washington, Friday Harbor Laboratories, University of Washington, 620 University Rd., Friday Harbor, WA 98250, USA, (MAK).

**Keywords:** Sexual dimorphism, Sexual selection, Geometric morphometrics, Gymnotiformes, Biomechanics

## Abstract

The evolution of sexual weaponry is thought to have marked effects on the underlying static allometry that builds them. These weapons can negatively affect organismal survivability by creating trade-offs between trait size and performance. Here we use three-dimensional geometric morphometrics to study the static allometry of two species of sexually dimorphic electric fishes (*Apteronotus rostratus* and *Compsaraia samueli*) in which mature males grow elongate jaws used in agonistic male-male interactions. We quantify jaw mechanical advantage between the sexes of both species to track changes in velocity and force transmission associated with the development of sexual weaponry. We find evidence for trade-offs between skull shape and mechanical advantage in *C. samueli,* where males with longer faces exhibit lower mechanical advantages, suggesting weaker bite forces. In contrast, males, and females of *A. rostratus* exhibit no difference in mechanical advantage associated with facial elongation. We hypothesize that differences in the functionality of the sexual weaponry between the two species may drive divergences in the allometric scaling of mechanical advantage.

## Background

Sexually-selected traits used as weapons in competition for resources, and ultimately access to mates, have evolved multiple times across the tree of life, and have produced a diversity of elaborate phenotypes [1]. These weapons are most often used to defend resources and settle conflicts between individuals through combat (armament) or display (ornament), and may also serve the additional purpose of providing honest signals to mates about viability [2, 3]. Sexual weaponry may also result in trade-offs between the size of a weapon and performance thus limiting the potential range of phenotypic disparity [4, 5]. Trade-offs also have marked effects on the underlying static allometries that build sexual weapons, such that differences in performance associated with increases in trait o size can influence the slope of a static allometry [6].

### Study System

Neotropical electric fishes (Gymnotiformes: Teleostei) represent an excellent case study for sexual weaponry. In this clade, fishes exhibit a wide diversity of skull shapes ranging from highly foreshortened skull shapes to highly elongate skull shapes [7, 8]. Amongst this diversity is an interesting pattern of sexual dimorphism, where males grow elongate snouts and oral jaws for use in agonistic jaw-locking battles, a trait that has evolved multiple times independently [9-11] (Figure S1; Movie 1).

Facial elongation of sexually dimorphic males presents an interesting case for the study of trade-offs in jaw biomechanics. The typical teleost jaw is an integrated system of levers and linkages that control the opening and closing of the jaws in feeding and other activities [12, 13]. There is an extensive literature that documents the biomechanical effects of changes in jaw lever lengths and the resulting functional consequences for performance [14-16]. The elongation of the snout and oral jaws in sexually dimorphic electric fishes may therefore result in trade-offs in jaw performance, as the lengths of jaw levers vary ontogenetically between the sexes.

Here we use three-dimensional geometric morphometrics to study the static allometry of sexually dimorphic phenotypes in two species of apteronotid electric fishes: *Apteronotus rostratus* and *Compsaraia samueli*, both of which exhibit independently-evolved craniofacial weaponry in males, and track ontogenetic trade-offs in mechanical advantage between the sexes in each species.

## Materials and Methods

### Specimen selection and preparation

For *Apteronotus* a total of 31 (12 male and 19 female) specimens were collected in the Chepo region of Panama (December-March 2016) using dip nets. To increase sample size, specimens collected in the field during this period were pooled with 26 museum specimens (five males, 21 females) from the Smithsonian Tropical Research Institute (STRI) collected from nearby (<20 km).

For *Compsaraia samueli*, a total of 49 (26 male and 23 female) specimens were collected from the area near Iquitos, Peru (August 2015-January 2017) using purse seines. Specimens of both species were dissected, and gonads inspected to determine sex (Table S1).

### Micro-CT scanning

We used micro-CT scanning to capture the osteological properties of individuals in three-dimensions. For *A. rostratus*, a size series of 30 specimens (49-212 mm TL) was scanned at the University of Texas, Austin (UT) using a custom-built scanner by North Star Imaging (NSI) at 180 kV, 114-115 uA and 19-49 μm voxel size. The remaining 27 specimens were scanned at the University of Washington Friday Harbor Labs (UW) Karl Liem Memorial Bio-Imaging Facility in conjunction with the “ScanAllFishes” project using a Bruker Skyscan 1173 at 70 kV, 114 uA and 28.2 μm voxel size. For *C. samueli*, a size series of 49 specimens (67-194 mm TL) were scanned at (UW) at 65-70 kV, 114-123 uA,24-35.7 μm voxel size, 1175-1200 ms exposure, and a CCD sensitivity of 2240 x 2240 pixels. All micro-CT scans of both species are freely available for download from Open Science Framework © at osf.io/m8tqe.

### Three-dimensional Geometric Morphometrics

To study the ontogenetic shape change of the neurocranium between sexes of the two species, we used three-dimensional geometric morphometrics. Micro-CT image stacks were imported into Stratovan Checkpoint© and converted to three-dimensional isosurfaces. Isosurfaces were digitized with 34 landmarks (LM)(13 bilaterally symmetrical) (Figure 1a-c:Table S2) and exported to *MorphoJ* [17] for subsequent statistical analysis.

**Figure 1.**
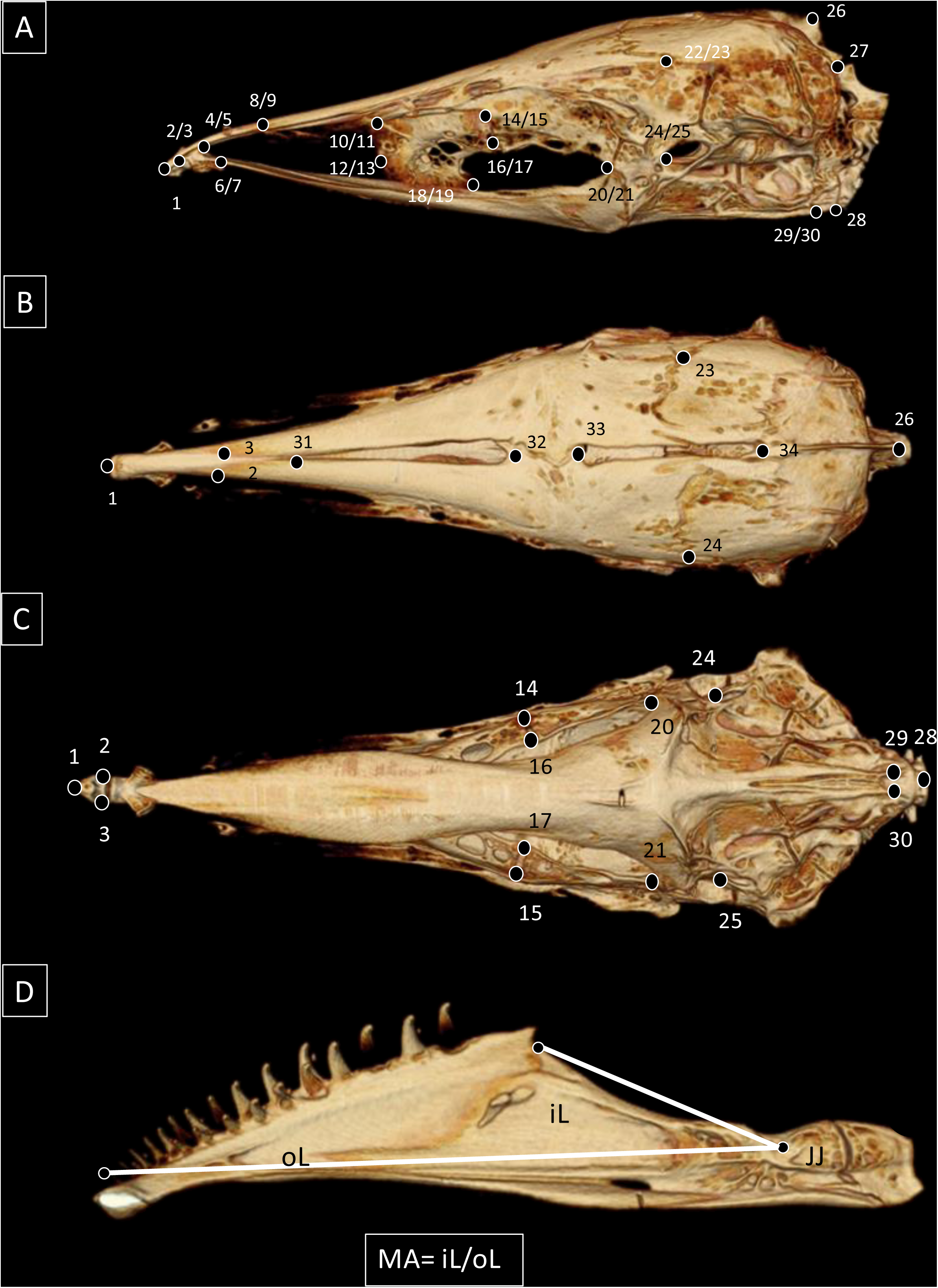
CT scans of *Apteronotus rostratus* (ANSP 200222) in lateral (A), dorsal (B), ventral (C), and mandibular views showing three-dimensional landmarks for 34 (13 bilaterally symmetrical) landmarks used for the geometric morphometric analysis of *Compsaraia samueli* and *Apteronotus rostratus* and in-lever (iL) and out-lever (oL) measurements taken from the jaw joint (JJ).

### Mechanical Advantage

We estimate velocity and force transmission of the lower jaw using closing mechanical advantage (MA). MA is quantified as the ratio between the closing in-lever distance (distance between the insertion of the adductor mandibulae muscle and the articulation of the jaw joint) and out-lever distance (distance between jaw joint and most anterior tooth) [13], such that MA= iL/oL (Figure 1d). To determine the precise area of muscle insertion on the lower jaw, one representative specimen of each species was dissected to reveal the underlying area of insertion. Here we study the ontogenetic scaling of log-transformed MA with log-transformed body-size and skull shape in male and female specimens of *A. rostratus* and *C. samueli* to test for differences in slopes between sexes in each species and identify potential performance trade-offs associated with facial elongation.

### Neurocranial Allometric Trajectories

To remove the effect of differential scaling and orientation, a full Procrustes superimposition was performed in *MorphoJ*. The Procrustes coordinates were then used to study the allometric relationship between skull shape and MA. Variation in allometric slopes between sexes was assessed in the R-package *Geomorph* [18] using the “procD.allometry” function. Allometric slopes are displayed using a predicted shape vs. MA regression [19].

### Video Recordings of Behavior

Agonistic interactions between *C. samueli* males were filmed at the “Amazon Tropicals” aquarium store in Iquitos, Peru. Individuals were collected by aquarium fishermen and housed in 40-gallon aquariums where they were filmed by the authors using a GoPro Hero 5© at 240 fps. Videos were rendered at 60 fps using Adobe Premier Pro Creative Cloud©.

## Results

### Mechanical Advantage in Apteronotus rostratus

No significant relationship was found between skull shape and MA in *A. rostratus* (Table 1; Figure 2a). Additionally, no significant interaction was found between MA and sex, indicating that males and females exhibit similar MA throughout the entirety of their ontogenies.

**Figure 2.**
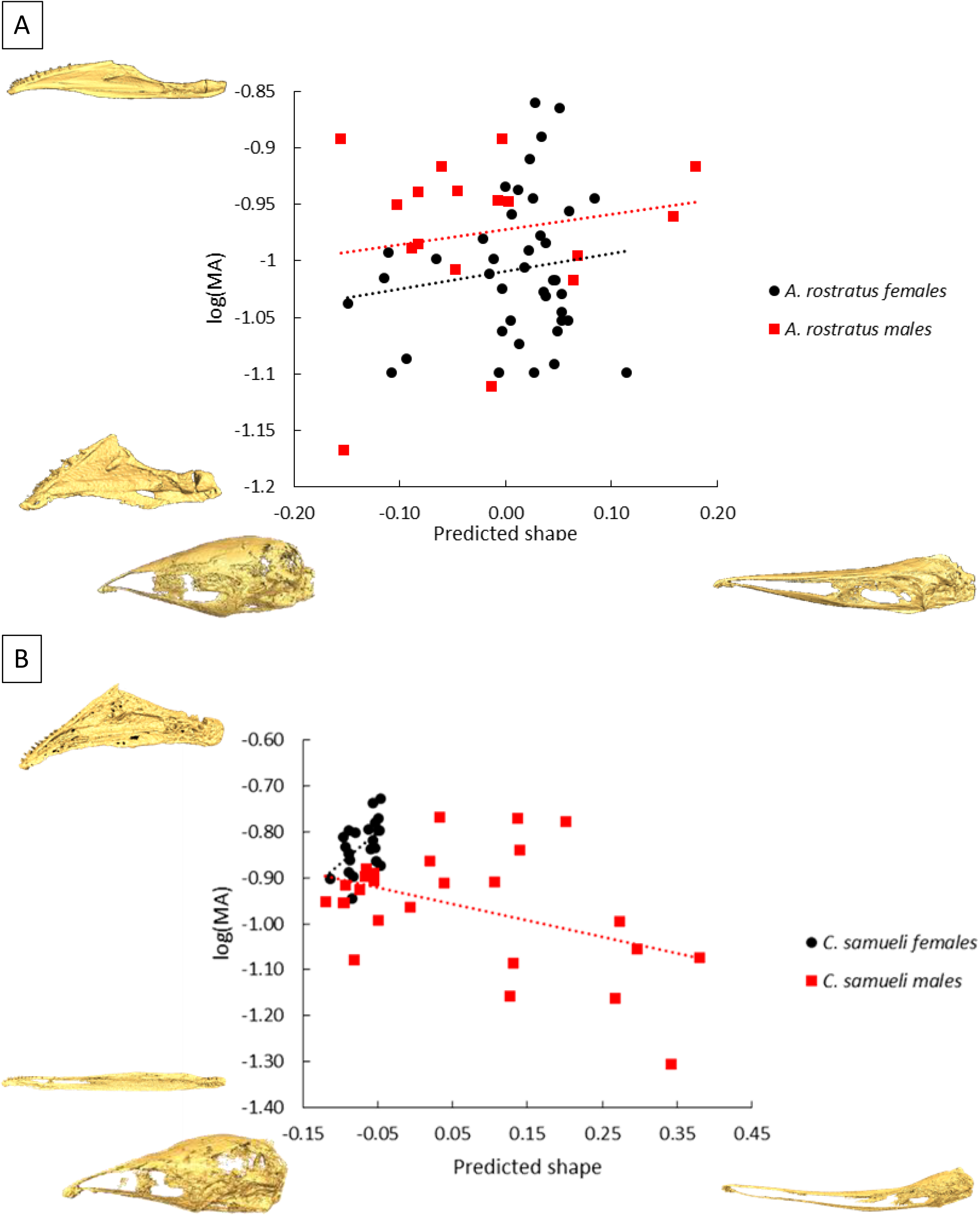
Ontogenetic trajectories of lower jaw mechanical advantage vs predicted skull shape in *Apteronotus rostratus* (A) and *Compsaraia samueli* (B).

**Table 1.** Analysis of variance for the effect of skull shape and sex on mechanical advantage for *Apteronotus rostratus* and *Compsaraia samueli*. Bold values indicate statistical significance (p= < 0.05).

| *A. rostratus* | Df | Sum Sq | Mean Sq | F value | Pr(>F) |
| --- | --- | --- | --- | --- | --- |
| shape | 1 | 0.003 | 0.003 | 0.595 | 0.444 |
| sex | 1 | 0.016 | 0.016 | 3.592 | 0.064 |
| shape:sex | 1 | 0.000 | 0.000 | 0.010 | 0.920 |
| Residuals | 45 | 0.238 | 0.004 |  |  |
| *C. samueli* | Df | Sum Sq | Mean Sq | F value | Pr(>F) |
| shape | 1 | 0.205 | 0.205 | 23.960 | **0.000** |
| sex | 1 | 0.065 | 0.065 | 7.590 | **0.008** |
| shape:sex | 1 | 0.030 | 0.030 | 3.484 | 0.068 |
| Residuals | 45 | 0.385 | 0.009 |  |  |

### Mechanical Advantage in Compsaraia samueli

A significant relationship was found between skull shape and MA where males with longer faces exhibit lower MA and females the opposite pattern (Table 1; Figure 2b). A significant effect of sex on MA was also recovered, suggesting differing MA between sexes.

## Discussion

### The performance cost of facial elongation differs among species

Notable scaling differences in MA were observed among species. In *C. samueli,* skull shape in males is negatively correlated with MA while females exhibit an inverse pattern. This reduction suggests that sexually dimorphic males have weaker jaw-closing forces than females, suggesting a trade-off in male cranial morphology, whereby males with elongate faces used in jaw-locking combat sacrifice more forceful biting commensurate with shorter jaws. This pattern is contrasted with *A. rostratus,* which exhibit no differences in MA between males and females.

### Why the long face?

Despite their exaggerated snout and jaws, male *C. samueli* have low MA jaws reaching as low as 27% (vs. 40% in the lowest female). Fittingly, observations of their fighting behavior (Movie 1) demonstrate that combat rarely results in extensive damage. This is a common finding among studies of animal weaponry where the function of an exaggerated weapon is greatly diminished and instead functions as an assessment tool for conspecifics [20]. There are several alternative explanations for low MA jaws in *C. samueli*: (1) lower, MA may be selected for in this taxon. This suggests that these jaws could be used as a more exclusionary weapon [21]. Observations show fighting *C. samueli* males facing each other head-on (Movie 1), without any obvious rolling or twisting, but instead pushing each other linearly in a contest more analogous to sumo-wrestling or tug-of-war, where opponents are pushed or pulled off-balance around a central arena.

(2) Hypermorphic jaws could reflect increasing ritualization of the structure, suggesting jaws are more ornamental and less useful as a functional weapon. Exaggerated features are typical of high-quality males, and serve as a clear signal to rivals that their competitor is robust, capable of defending a resource, and not worth fighting with. Facial elongation also results in the increase in absolute body-length of these and may be made obvious through electric organ discharge by increasing the distance between dipoles.

## Conclusions

Differences in the mechanical advantage of convergent sexual weaponry between these two species reflects a functional gradient between armament and ornamentation that is seen across other taxa. In *A. rostratus*, males retain a fully functional lower jaw throughout ontogenetic skull elongation. The opposite pattern is observed in *C. samueli* males where the mechanical advantage of the lower jaw is dramatically reduced resulting in a less functional weapon but perhaps a more appealing or ritualized ornament for conspecific signaling.

## Acknowledgments

We thank Brandon Waltz, Eric Hilton, Andrew Simons and William Crampton for discussions about sexual dimorphism Adam Summers and #scanAllFishes project allowed us to scan specimens for free.

## Ethics

Fieldwork: IACUC 2016-0325-2019

## Data Accessibility

Shape data with script available on Dryad: doi:10.5061/dryad.mh911

## Author Contributions

KME wrote the manuscript, collected specimens and took shape measurements. MJB assisted with specimen collection and scanning. MAK assisted with data interpretation and scanning. KLF filmed specimens in Peru and assisted in CT specimens. JSA reviewed the manuscript and assisted in data interpretation.

---

**Movie 1.** Agonistic jaw-locking behavior between two captive male *Compsaraia samueli* specimens.

